# A test of the hypothesis that variable mutation rates create signals that have previously been interpreted as evidence of archaic introgression into humans

**DOI:** 10.1101/2020.12.23.424213

**Authors:** William Amos

**Author notes:** **Author and corresponding author:** William Amos.

## Abstract

It is widely accepted that non-African humans carry 1-2% Neanderthal DNA due to historical inter-breeding. However, inferences about introgression rely on a critical assumption that mutation rate is constant and that back-mutations are too rare to be important. Both these assumptions have been challenged, and recent evidence points towards an alternative model where signals interpreted as introgression are driven mainly by higher mutation rates in Africa. In this model, non-Africans appear closer to archaics not because they harbour introgressed fragments but because Africans have diverged more. Here I test this idea by using the density of rare, human-specific variants (RHSVs) as a proxy for recent mutation rate. I find that sites that contribute most to the signal interpreted as introgression tend to occur in tightly defined regions spanning only a few hundred bases in which mutation rate differs greatly between the two human populations being compared. Mutation rate is invariably higher in the population into which introgression is *not* inferred. I confirmed that RHSV density reflects mutation rate by conducting a parallel analysis looking at the density of RHSVs around sites with three alleles, an independent class of site that also requires recurrent mutations to form. Near-identical peaks in RHSV density are found, suggesting a common cause. Similarly, coalescent simulations confirm that, with constant mutation rate, introgressed fragments do not occur preferentially in regions with a high density of rare, human-specific variants. Together, these observations are difficult to reconcile with a model where excess base-sharing is driven by archaic legacies but instead provide support for a higher mutation rate inside Africa driving increased divergence from the ancestral human state.

## Introduction

Around a decade ago came the revelation that humans and Neanderthals inter-bred, leaving a persisting legacy of around 2% in non-Africans (Green *et al.* 2010; Prufer *et al.* 2014; Sankararaman *et al.* 2016; Vernot *et al.* 2016). This legacy was originally inferred by D, a measure of asymmetric allele sharing between the Neanderthal genomes and one of a pair of human populations (Martin *et al.* 2015). While a number of other measures have subsequently been proposed (Meyer *et al.* 2012; Patterson *et al.* 2012; Mallick *et al.* 2016; Peter 2016), D is simple, transparent, has a strong analytical basis and still widely used. The initial revelation has generated much interest in the idea of inter-breeding as a crucial element in hominin evolution, and led to revelations about the way introgressed fragments may have facilitated human adaptation (Huerta-Sanchez *et al.* 2014; Gittelman *et al.* 2016), and complicated patterns of mixing (Slon *et al.* 2018), involving Denisovans (Meyer *et al.* 2012; Qin and Stoneking 2015; Dannemann *et al.* 2016) and perhaps other, as yet undiscovered archaics.

The introgression hypothesis is not without issue. Haploid regions of the genome such as the Y chromosome, mitochondrial DNA and to some extent the X chromosome appear not to have introgressed (Sankararaman *et al.* 2014; Delser *et al.* 2017). This may have been prevented by natural selection (Dannemann *et al.* 2016), though it remains unclear whether the loss of fitness needed to prevent any introgression of these regions would still have allowed hybrid lineages to persist and thrive. More recently, the signals interpreted as introgression have been shown to be dominated by heterozygous sites in Africans (Amos 2020), the exact opposite of the expected pattern where introgressed fragments, which are expected mostly to be rare, and should therefore occur mainly in the heterozygous state outside Africa. This observation highlights a potential problem with the inference of introgression. While it is assumed that non-Africans are more similar to archaics because their genomes harbour archaic fragments, an alternative possibility is that African sequences are less similar to archaics due to a higher mutation rate. Studies reporting introgression universally assume that mutation rate is constant (Green *et al.* 2010; Patterson *et al.* 2012) and, as a result, this alternative model has not previously been considered.

I have recently explored this problem by asking about the types of site that contribute to D and the sequence context in which they lie (Amos in review). Surprisingly, D is dominated by sites where the putative introgressed archaic allele is fixed in populations outside Africa rather than, as predicted by introgression, rare. Moreover, there is a strong tendency for sites that contribute most strongly to D to occur in regions where flanking heterozygosity is much higher in one human population compared with the other. Specifically, in any given genomic region, the magnitude and sign of D persistently imply introgression into the population with lower heterozygosity. This trend holds even where genome-wide D indicates introgression in the opposite direction.

The signal previously interpreted as indicating archaic legacies in humans thus appears to be dominated by sites where the putative archaic allele is fixed in non-Africans (Amos in review), by sites that that tend to be heterozygous in Africans (Amos 2020) and by sites that occurs preferentially in genomic regions where flanking heterozygosity is unusually higher in Africans compared with non-Africans (Amos in review). Considered together, these trends appear consistent with a speculative new model where heterozygosity modulates mutation rate such that the large loss of heterozygosity ‘out of Africa’ led to a general mutation rate reduction in non-Africans (Amos 2011; Amos 2013). Here, the higher African mutation rate acts to drive increased divergence from archaics rather than introgression acting to reduce divergence between Neanderthals and non-Africans. This model is attractive because it explains a number of observations that a simple introgression model struggles to account for: the west to east increase in inferred legacy across Eurasia (Wall *et al.* 2013), which parallels the way heterozygosity declines with distance from Africa (Prugnolle *et al.* 2005); the apparent lack of introgression of haploid / semi-haploid where an effect of heterozygosity would be reduced or lacking; the fact that the signal is carried by heterozygous sites inside rather than outside Africa (Amos 2020); and, most directly, the way flanking sequence heterozygosity *difference* predicts D.

Here I attempt a direct test of the idea that in humans, D is driven by variation in mutation rate. For this, I systematically identify bases that contribute to signals interpreted as introgression and attempt to quantify the mutation rate context in which each such base lies. I find evidence that bases contributing to the signals interpreted as introgression lie in very narrow genomic regions where mutation rate is appreciably higher in the population where introgression is not inferred.

## Results

### a) Quantifying the impact of local mutation rate variation

D is dominated by sites where the Neanderthal allele is *fixed* in non-Africans, suggesting strongly that D is driven mainly by back-mutations from the common BBBA state, one substitution separating hominins from the chimpanzee. The genome-wide average mutation rate of a little over 10^−8^ (Besenbacher *et al.* 2016) is too low to generate more than a handful of back-mutations. Consequently, for back-mutations to contribute appreciably to D, D-informative bases (DIBs) must lie preferentially in mutation hotspots. To test this prediction by determining mutation rate directly and accurately across the genome is prohibitively costly. However, on the assumption that most substitutions are neutral, a reasonable estimate of local mutation rate can be obtained by counting the numbers of sites carrying rare, human-specific derived variants (RHSVs), defined as rare human variants not present in the Neanderthal, Denisovan or chimpanzee.

To explore the mutational context of putative introgressed bases, I systematically identified all DIBs. For each such base I scanned a pair of 1Kb genomic windows either side for RHSVs, counting the number in each population. I defined RHSVs as bases present in more than one copy but fewer than 200 copies across the entire 5008 alleles called at each site in the 1000 genomes Phase 3 data. Singletons were excluded in case they are enriched for sequencing errors. The 200 copy upper limit was chosen as a balance between maximising the number of variants included while at the same time minimising the number of variant that predate ‘out of Africa’, but was otherwise arbitrary. The frequency of RHSVs reflects both local mutation rate and local time to most recent common ancestor (TMRCA). Variation in TMRCA along a chromosome will create noise, but should not drive systematic trends (an assumption that was confirmed by simulations, see below). Therefore, at each site I used the difference in RHSV number between populations as a surrogate measure of difference in local mutation rate. For every level of difference, I then used the associated DIBs to calculate the expected number of ABBAs and BABAs and hence D (shown diagrammatically in Figure 1).

**Figure 1.**
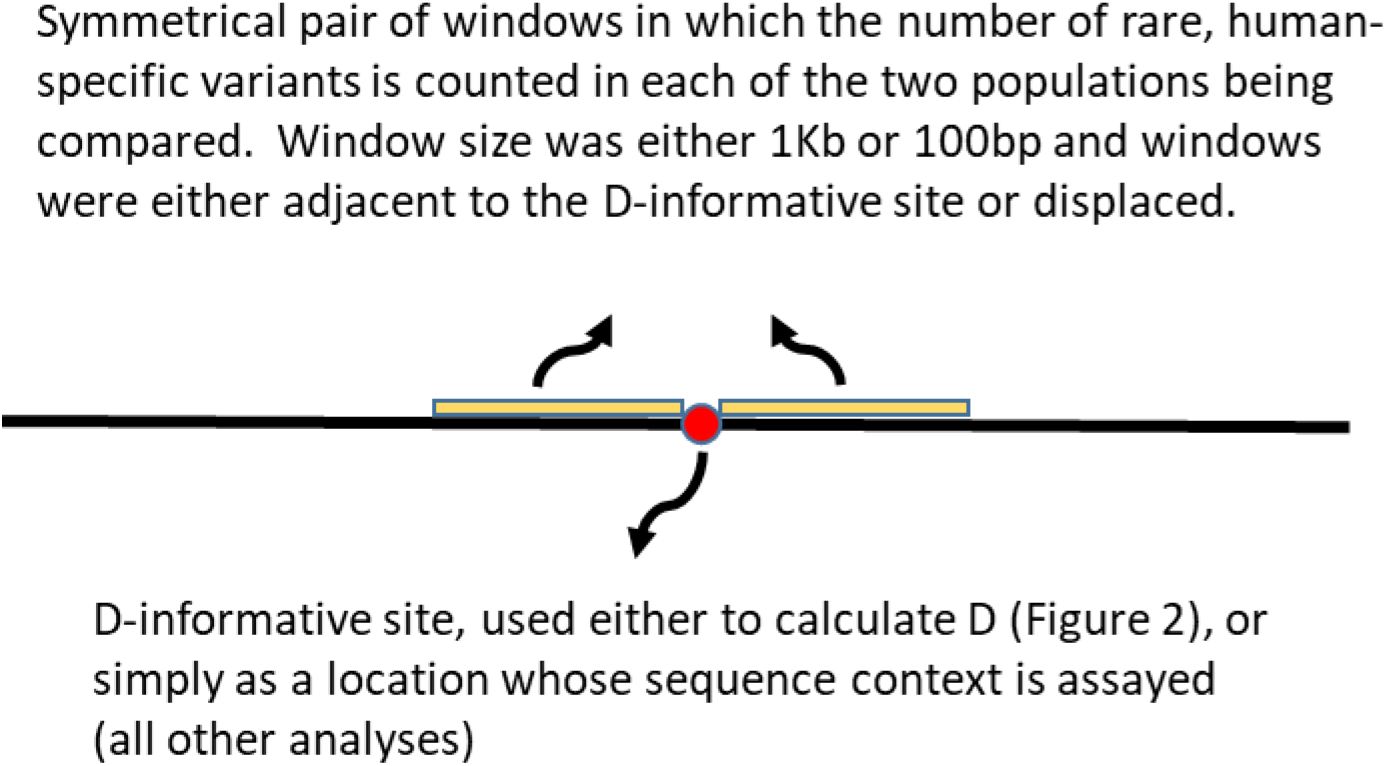
Analytical design. To establish the mutational context of any focal base of interest (red circle), such as a D-informative site (as illustrated), pairs of symmetrically placed, 1Kb counting windows (yellow) were placed either side. Other focal base types explored were bases carry three alleles in humans and bases carrying human-specific derived alleles. The counting windows were moved outwards in kilobase steps to explore the dependence of any pattern on distance from the focal site, or divided into 20 sub-windows for a fine-scale analysis.

To visualise the extent of spread of any trend, I used pairs of 1Kb windows placed symmetrically either side of each DIB, displaced by some number of kilobases (range 0 to 20). Results were averaged across all like regional comparisons (i.e. all Europe-Africa comparisons were average to produce a single EUR-AFR set of values) and are presented as four comparisons that illustrate the major trends within Africa, outside Africa and between African and non-African populations: EUR-AFR, EAS-AFR, AFR-AFR and EUR-EAS (Figure 2). In the two Eurasia – Africa comparisons (top panels) most data lie to the left of zero on the X-axis, confirming the larger numbers of RHSVs in Africa, and show a strong negative slope indicating that D is larger the greater the excess number of RHSVs in Africa. Extreme values are based on many fewer sites and are therefore more variable. Also noticeable is the near perfect ordering by window displacement: the steepest lines are linked to windows immediately flanking each focal base (displacement = 0, black), and slopes get progressively shallower as the counting windows are moved further away from the DIB, becoming almost flat when displaced by 20Kb (purple). Comparisons within Africa (bottom left panel) are both more symmetrical and less variable, likely due to a combination of the larger number of informative sites and their simpler demographic histories that lack the ‘out of Africa’ bottleneck. Finally, comparisons outside Africa, illustrated by EUR-EAS, show flatter, more variable patterns across a much narrower range of RHSV differences.

**Figure 2.**
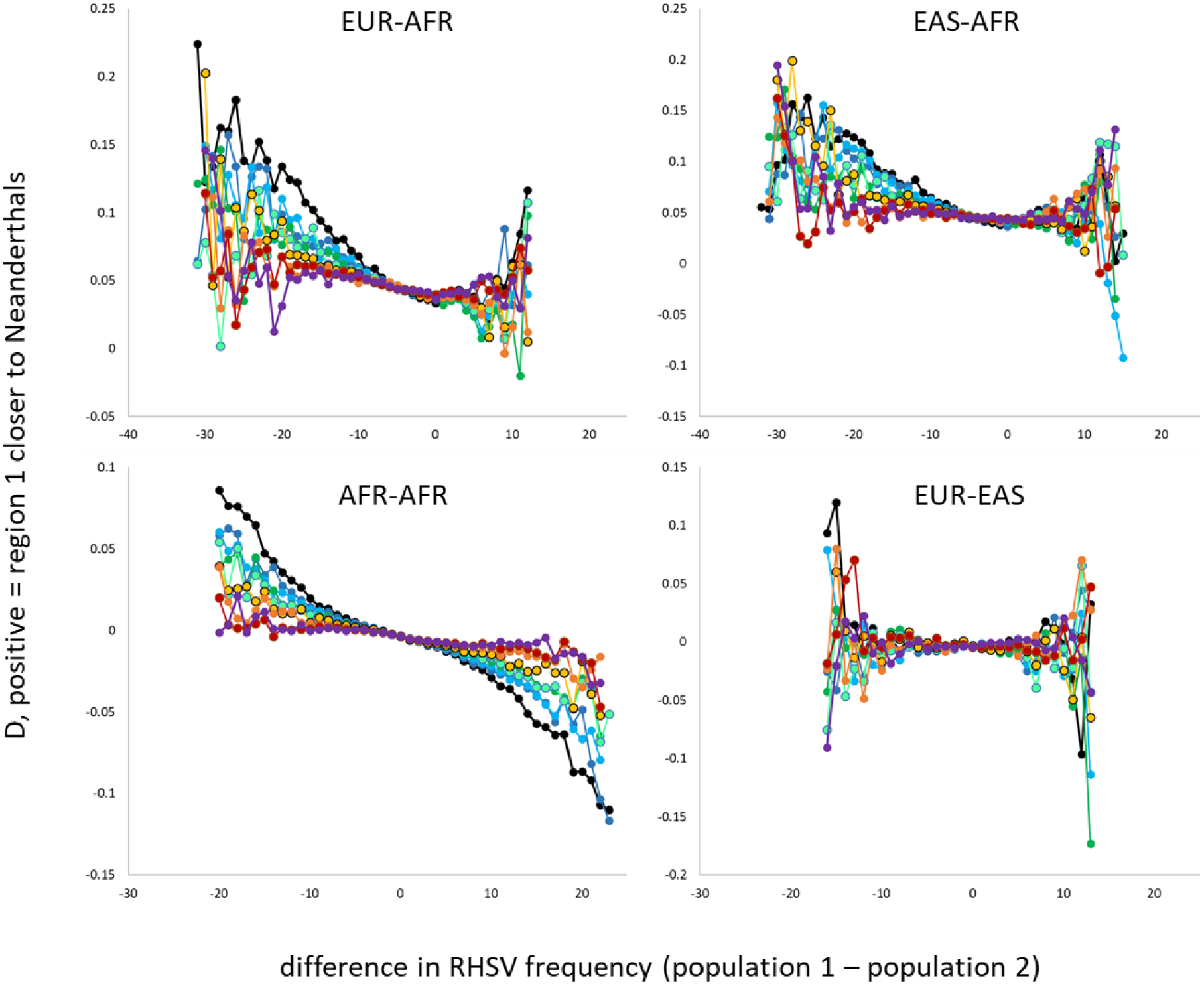
Dependence of D on relative flanking sequence mutation rate. In every pairwise population comparison, every D-informative site (DIS) was identified and treated as the focal base. 1Kb windows were then used to count flanking rare, human-specific variants in each population separately, the two counts being combined as a difference in number, with counts in population two subtracted from counts in population one (X-axis). All DISs with the same count difference were then used to calculate a value of D (Y-axis). The process was repeated, with counting windows starting at 0 (i.e. adjacent to the focal base, black) and moved progressively further out (1, 2, 3, 4, 5, 10, 15, 20Kb, respectively colours moving through dark blue, light blue, green, light green, yellow, orange, red and purple). Data are presented as averages across all equivalent population comparisons that represent a given regional comparison, i.e. all profiles with any European population in position one and any African population in position two were averaged to produce one EUR-AFR comparison. Four different comparisons are presented to illustrate the classic Africa – non-Africa case (EUR-AFR and EAS-AFR), within Africa (AFR-AFR) and outside Africa (EUR-EAS).

### b) Exploring the average mutational context of sites that contribute to D

DISs appear to sit preferentially in regions where heterozygosity (Amos in review) and mutation rate (see above) are higher in the population into which introgression is *not* inferred. Moreover, a large majority of DISs involve sites where one of the two alleles is fixed in one of the two human populations. For a fine-scale view of the mutational environment in which an average DIS sits, I divided the 1Kb counting windows into 20 sub-windows, each of just 100bp long. DISs were classified into four types according to whether the A or B allele was fixed in one population and whether the Neanderthal base was likely ancestral (Neanderthal and Denisovan carry the same base) or derived (Neanderthal and Denisovan bases differ). For each population / region comparison, I calculated a standardised measure of difference in RHSV number:

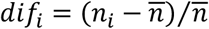

where *n*_*i*_ is the difference in RHSV counts in sub-window *i* between the two populations, calculated as the number of RHSVs in the population where the DIS carries both A and B minus the number of RHSVs in the population where either A or B is fixed. Representative plots are presented in Figure 3, illustrated by comparisons in which A and B are segregating in either GBR or ESN carry the fixed allele, average across all instances in which A or B is fixed in the comparator population. Positive standardised RHSV differences indicate a higher than average frequency of RHSVs in the population where A and B are both present. All four DIS types sit in the middle of a clear central peak and this peak is much stronger for DISs where the putative Neanderthal B allele is both likely ancestral and fixed in the focal population (GBR / ESN). Perhaps tellingly, this is also the type of DIS that accounts for a large majority of the signal captured by D.

**Figure 3.**
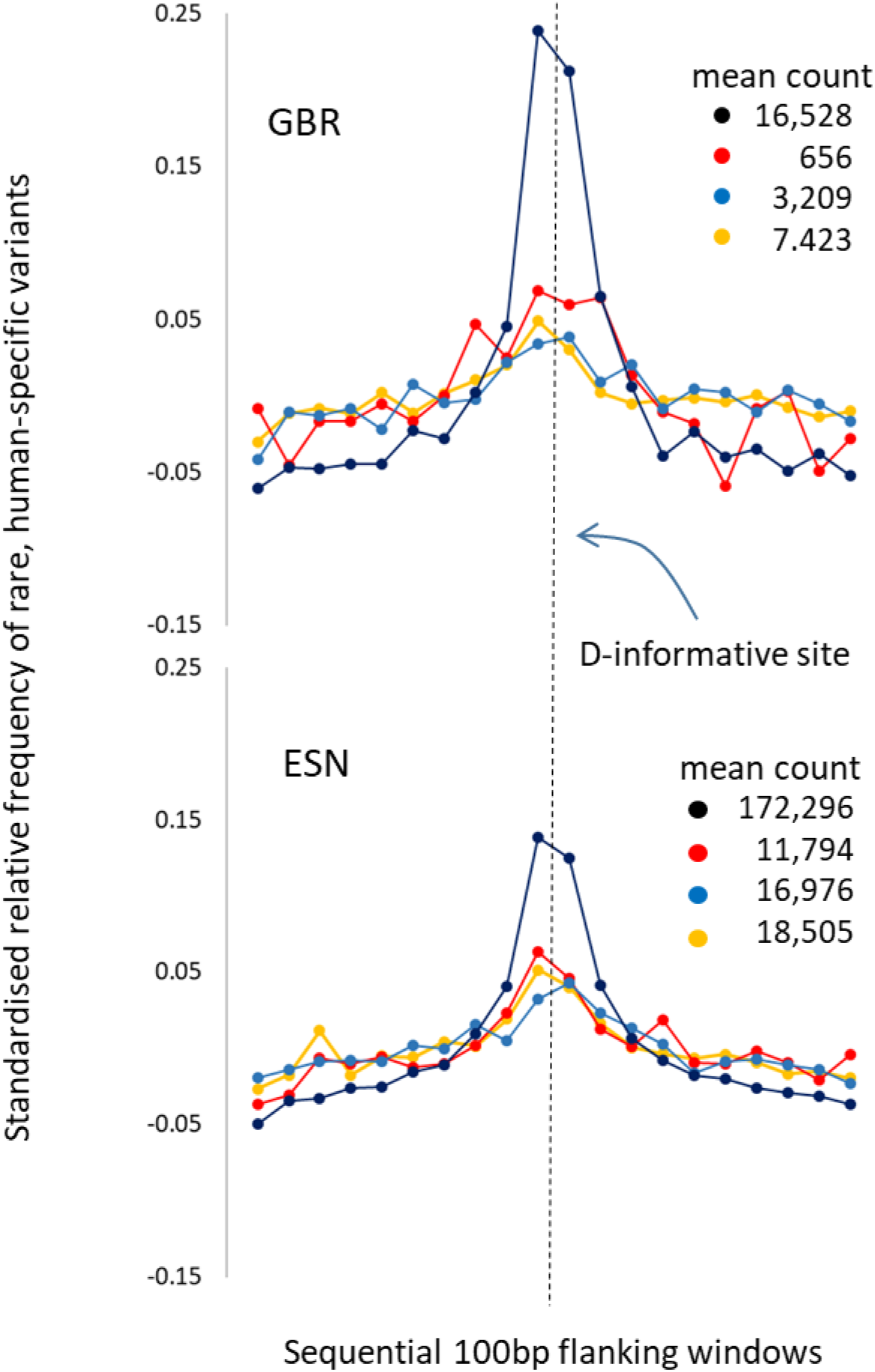
Variation in mutation rate in relation to distance from D-informative sites. All D-informative sites (DISs) were identified where one of the two bases was fixed in one of the two populations being compared. Each such base was classified according to: (a) whether the archaic base was likely ancestral (Neanderthal and Denisovan bases the same) or derived (Neanderthal and Denisovan differ); and (b) whether the fixed allele was shared with Neanderthals (i.e. B) or not (i.e. A), giving four different classes. One non-African population (GBR) and one African population (ESN) were selected at random. Total RHSV counts were generated for 1Kb windows placed either side of the DIS and sub-divided into 20 x 100bp sub-windows. I was interested in the possibility that back-mutations occur preferentially in regions of higher mutability, so counted RHSVs in the population where both A and B were present, summing across instances where the other population was fixed. The resulting counts were normalised relative to each other (see methods) and plotted according to the class of DIS: black = Neanderthal allele is fixed and ancestral; blue = non-Neanderthal allele is fixed, Neanderthal allele is ancestral; red = Neanderthal allele is fixed and derived; yellow = non-Neanderthal allele is fixed, Neanderthal allele is derived. For reference, the average number of RHSVs (and hence, approximately) the average number of sites contributing to D are also given, emphasising the dominant contribution of ancestral Neanderthal alleles fixed in one population (black).

The above analysis shows that relative recent mutation rate, as indicated by the frequencies of RHSVs, exhibits local peaks around sites that contribute to D where either A or B is fixed in one of the two populations, being higher in the population where both alleles are segregating. This pattern is consistent with the idea that many DISs feature fixed ancestral alleles that have subsequently back-mutated in highly localised mutation hotspots. For a broader context, I next attempted to compare the density of RHSVs near DISs with the overall genome-wide rate. This is difficult to conduct fairly because sequences flanking DISs are likely to have been called in all taxa, while many randomly-sited windows may fall in regions where one or more taxa have not been sequenced to sufficient depth. There is also the ambiguity that gaps in vcf files can represent either invariant sequences or missing data. To circumvent this problem, I ignored windows containing zero RHSVs. I then standardised the frequencies of windows carrying multiple RHSVs relative to the number of windows containing just a single RHSV. Plots comparing regional average values for the general background rate (black), 100bp windows displaced by 1Kb from a DIS (red) and 100bp windows immediately adjacent to a DIS are presented in Figure 4. Relative frequency is always in the order adjacent window (purple, highest), displaced window (middle, red) and background window (lowest, black), indicating that 100bp windows near to DISs are consistently more likely to contain multiple RHSVs. Moreover, the gaps between the three window types increase with the number of RHSVs in a window, reaching almost 100 times the relative background frequency for windows containing nine RHSVs in Africa. Thus, DISs lie in regions that both locally, on a scale of 100bp, and genome-wide have an unusually high density of RHSVs and, by implication, a higher recent mutation rate.

**Figure 4.**
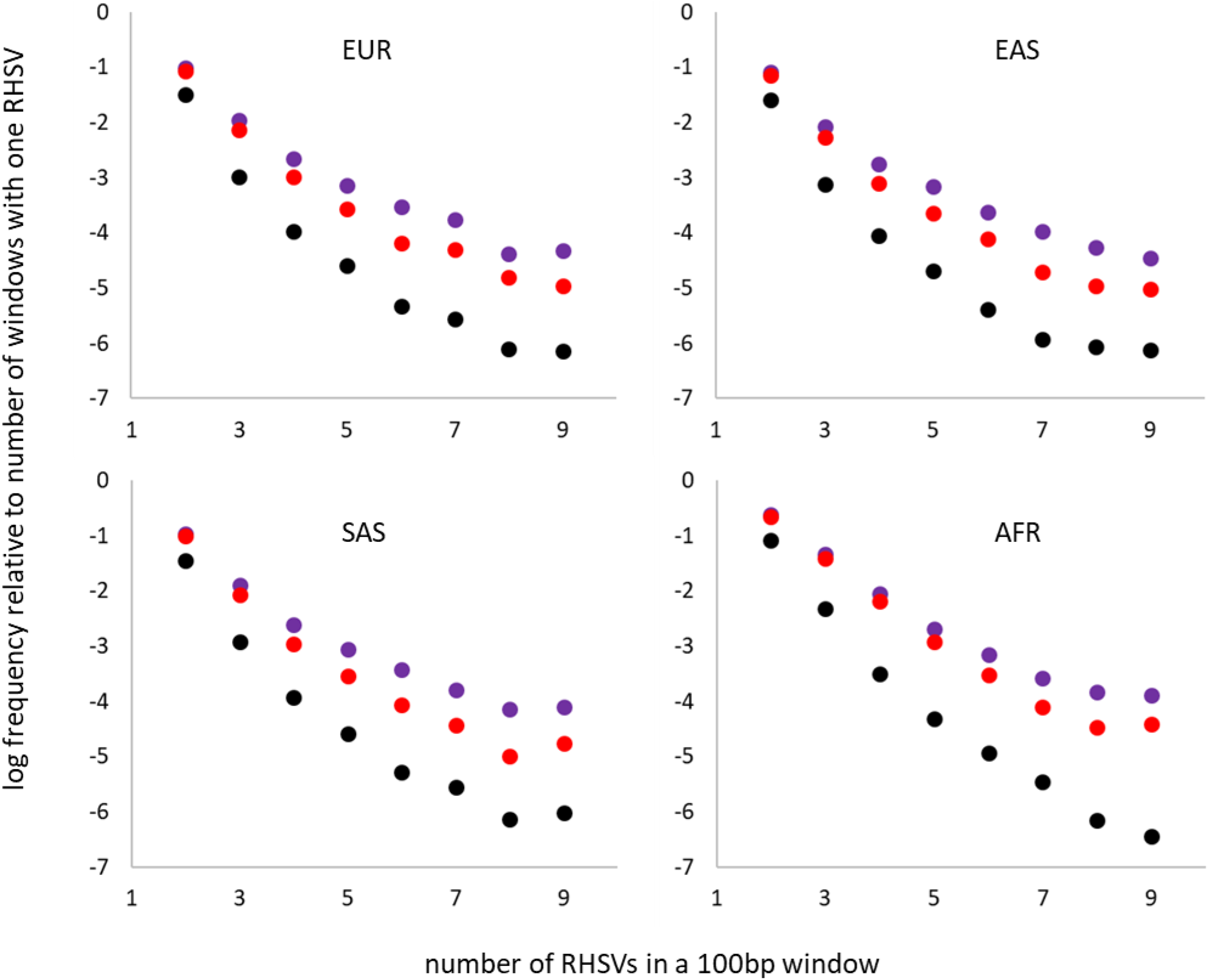
Mutation rate around D-informative sites relative to the background rate. The genome was divided into non-overlapping 100bp windows in which the numbers of rare, human-specific variants (RHSVs) were counted. Counts above nine were extremely rare and combined with the nine class. Since zero counts are difficult to interpret, often arising through missing data, counts between two and nine were standardised by dividing by the number of windows with one RHSV and taking the log (black). The process was repeated for 100bp windows adjacent to D-informative sites (DISs, purple) and for 100bp windows lying between 900bp and 1Kb displaced from DISs (red). The relative frequency of windows carrying multiple RHSVs increases predictably in order genome-wide average, near a DIS, adjacent to a DIS. Regional averages are presented for four main geographic regions: Europe, East Asia, South Asia and Africa. The highly admixed American populations give similar patterns but were omitted.

### c) Do excess rare variants reflect excess mutation rate?

To clarify the meaning of differences in RHSV counts, I sought a different class of site that have nothing to do with classical D yet should still reflect locally higher mutation rates. Across the 1000 genomes data, 257,000 sites carry three different alleles in humans (Amos 2020), vastly more can reasonably be attributed to sequencing errors in this highly-curated data set. Sites carrying three alleles (‘triples’) require two mutations at the same site, usually one transition and one transversion, and can be thought of as ‘visible’ back-mutations. As such, the mutational context of triples should be similar to the mutational context of sites where back-mutations create ABBAs and BABAs from the common BBBA state. To test this prediction, I conducted an exhaustive search for autosomal triples. At each one, I identified the rarest of the three alleles, presumed to be descended from the most recent mutation. In populations in which this rarest allele was present, I then counted RHSVs in the immediately flanking region, using the same 20 sub-window approach used above. All 26 populations yield very similar profiles and these are illustrated as standardised regional averages (Figure 5). Europe, East Asia and South Asia share near-identical profiles. The American peak is somewhat lower, possibly because in these populations many rare variants arise due to extensive admixture rather than mutations within that population. The lowest peak is in Africa and this likely reflects the substantially larger number of rare variants in these populations acting to inflate the mean density of RHSVs which in turn reduces the size of any peak following standardisation.

**Figure 5.**
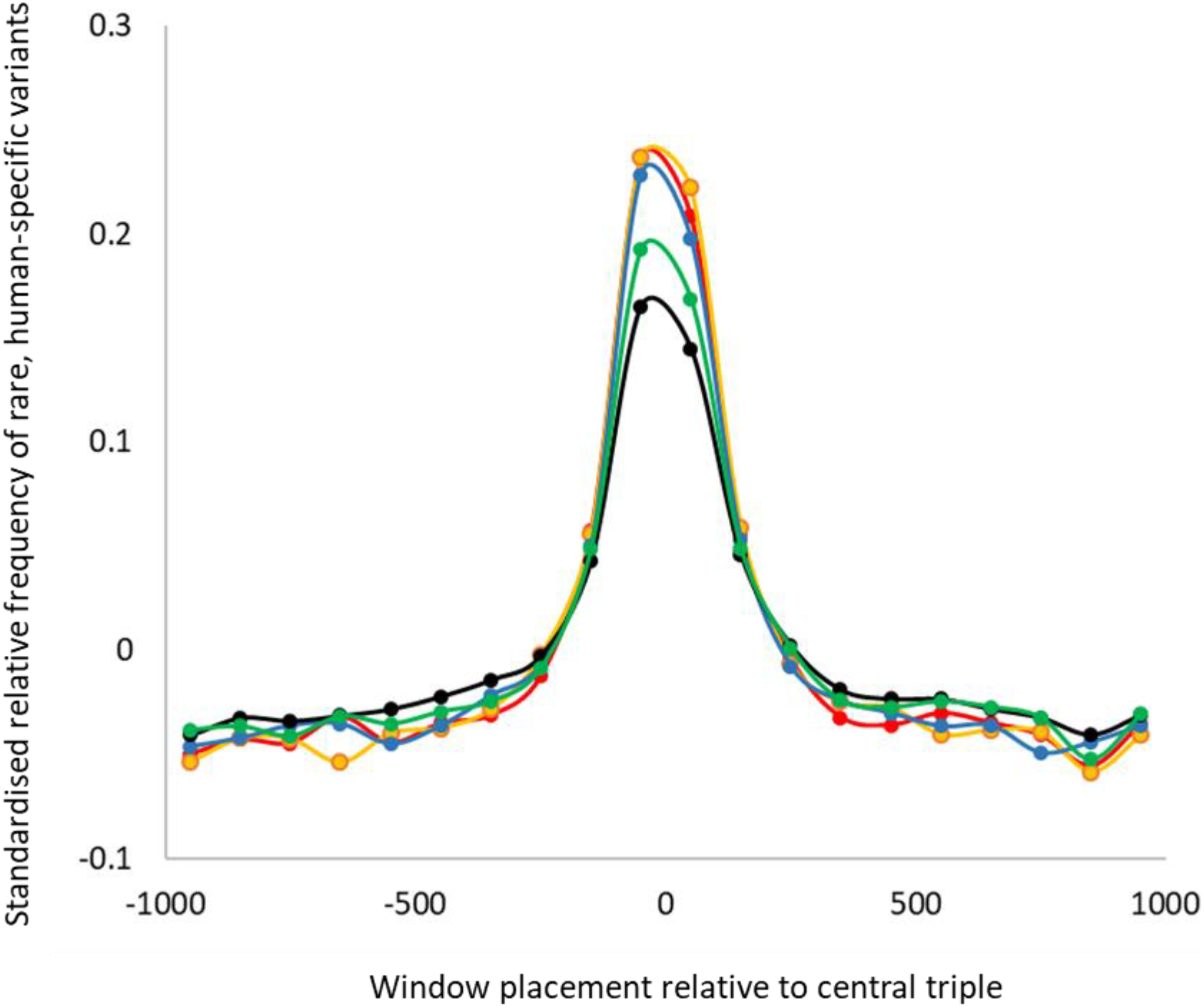
Verifying the meaning of high densities of rare variants. If mutation hotspots are genuinely responsible for generating D-informative sites (DISs) via back-mutations, similar patterns should be found surrounding other classes of variant that also require high mutation rates. I therefore sought all sites carrying three alleles, identified all instances where the rarest of the three alleles was present in more than one copy and fewer than 200 copies. I every population where such a base occurred, I counted recent, human-specific variants (RHSVs) in 20 x 100bp windows flanking but excluding the three-allele site. Standardised counts are presented as regional averages for Africa (black); Europe (red); East Asia (yellow); South Asia (blue); America (green).

### d) Coalescent simulations

To discover whether the observed patterns are consistent with a simple model based on introgression, I conducted a series of coalescent simulations. Four scenarios were explored: with and without Neanderthal introgression into the non-African lineage and with D calculated using either putative derived or putative ancestral archaic alleles. In each instance, I sought DISs and counted the numbers of RHSVs in both populations using a pair of adjacent windows each approximating 5Kb. This design replicates the analysis underpinning Figure 2 and results are presented in Figure 6. As expected, at the extremes of the distribution, scatter is much greater. However, within the region of each graph where ABBA and BABA counts exceed 10, all profiles are essentially flat and follow intuitive expectations. Within Africa (black) and outside Africa (red), D is consistently zero across all X-axis values. In the comparison between simulated Africans and non-Africans (open circles), the profiles are again flat, but are now positive, being larger for the derived than for putative ancestral alleles, reflecting the fact that putative derived alleles will include a lower proportion of ABBAs and BABAs generated by incomplete lineage sorting compared with putative introgressed ancestral archaic alleles. None of these simulations revealed any appreciable dependence of D on difference in the number of flanking RHSVs.

**Figure 6.**
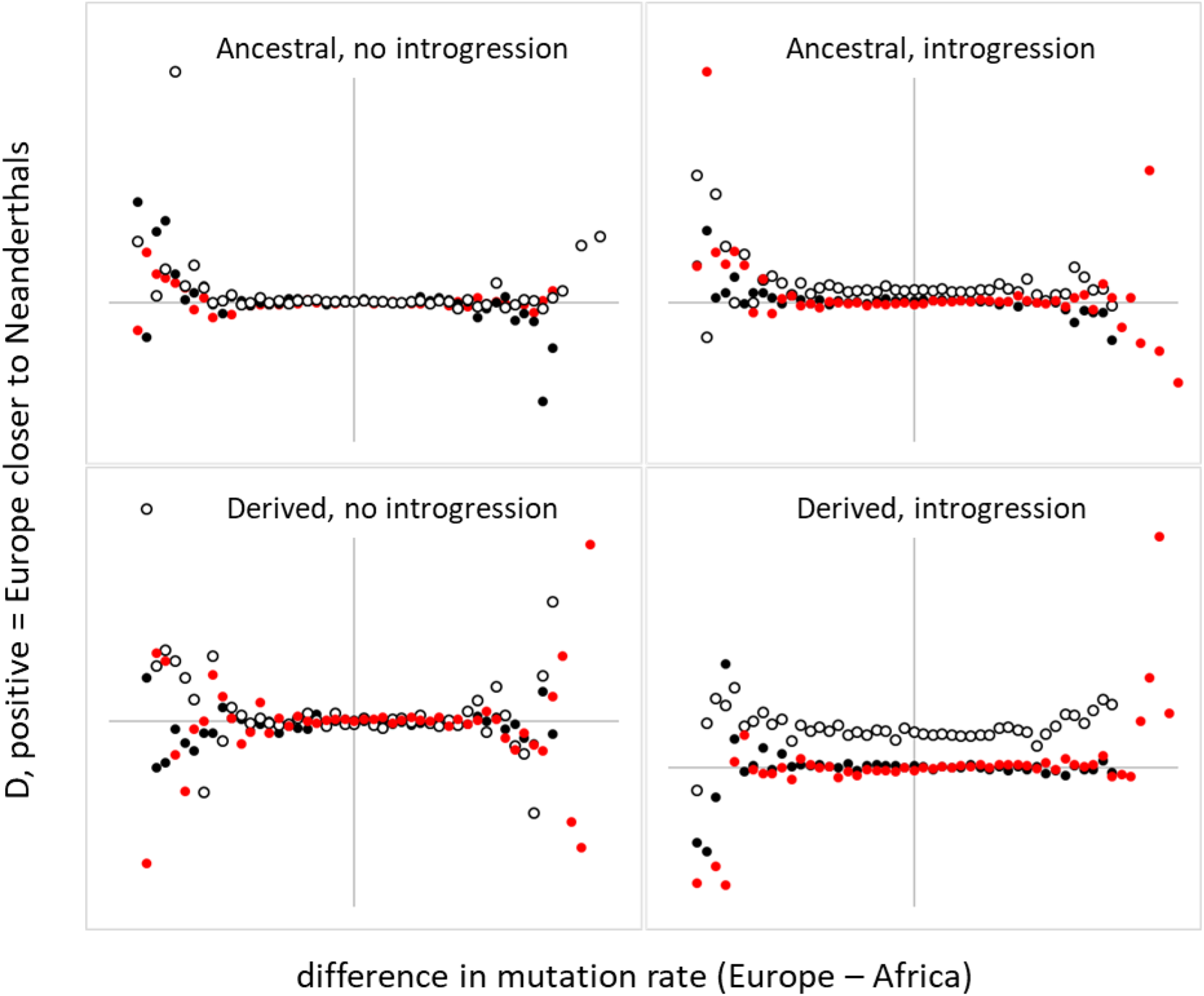
Simulated relationship between introgression and local mutation rate. Coalescent simulations were conducted in ms to produce a pair of ‘European’ populations, a pair of African populations, the Neanderthal and the chimpanzee. At each D-informative site I calculated the difference in number of rare, human-specific variants (RHSVs) in a pair of XKb windows immediately flanking but excluding the site. As in Figure 2, difference in RHSV number was used to define a series of bins, each of which yields a value for D, summing ABBA and BABA counts across all such similar sites. The analysis was repeated with and without simulated introgression, for putative ancestral and derived Neanderthal alleles, and for the three possible comparisons Europe-Europe (black), Africa-Africa (red) and Europe-Africa (white). As expected, D~0 when introgression is absent and within each region, becoming positive in Europe-Africa comparisons, with larger values when the Neanderthal allele is derived rather than likely ancestral.

## Discussion

Here I use the frequency of rare, human-specific variants as an indicator of mutation rate and show that sites contributing to D occur preferentially in what appear to be mutation hotspots. D is predictably larger in genomic regions when there is a large difference in mutation rate between populations. Refining this pattern reveals a sphere of influence only around 500bp wide. None of these trends are replicated coalescent simulations that assume an infinite sites mutation model.

Coalescent simulations confirm the intuitive expectation that the frequency of rare, human-specific variants, RHSVs, should be essentially unaffected by, and uncorrelated with, the presence of introgressed fragments. Thus, introgressed fragments are generally rare, so will have a negligible impact on the frequencies of human-specific alleles. Equally, by definition, introgressed fragments do not carry human-specific variants (strictly, the rate would be low rather than zero due to occasional, rare, derived Neanderthal alleles not present in the non-human genomes). Lastly, introgressed fragments are estimated to average around 50Kb in length, which is far too large to drive patterning on a scale of a kilobase or less. Despite this, a strong relationship exists between D and the frequency of RHSVs in the immediately flanking sequences. This relationship is most pronounced in African – non-African population comparisons and comparisons within Africa. Specifically, sites that contribute to large positive D values lie preferentially in regions where RHSVs are much commoner in the population where introgression is *not* inferred. Such a pattern is difficult to explain using a simple model based on introgression but it is consistent with the alternative model based on locally high mutation rates driving the formation of ABBAs and BABAs by back-mutations from the common BBBA state. Moreover, the back-mutation model also predicts the observation that D is dominated by sites where putative introgressed archaic alleles are ancestral rather than derived within Neanderthals, and hence reflects the ancestral human state rather than reflecting a specifically Neanderthal origin.

A very large majority of novel variants are lost by drift soon after they are formed (Kimura 1983; Mathieson and Mcvean 2014). Consequently, most rare variants have a relatively recent origin. It has been estimated that alleles present in only two copies in the 1000 genomes data are on average of the order of 200 generations old (Mathieson and Mcvean 2014) but average age outside Africa is around half that within Africa. This difference reflects a complicated relationship between the allele frequency spectrum in a population, that population’s history and stochastic variation in time to most recent common ancestor along a chromosome. Consequently, while the local density of rare, human-specific variants provides a reasonable proxy for local mutation rate, the measure is far from perfect, and its interpretation is more robust when it is used as a relative measure, for example by comparing values in a given population for series of windows along a chromosome. Comparisons between populations need to be treated with more caution.

I attempted to verify that the local density of RHSVs can be used as a proxy for mutation rate by focusing on sites in humans that carry three alleles. Such sites appear far too common to be explained by sequencing errors and require at least two mutations at the same site (Hodgkinson and Eyre-walker 2010; Amos 2020). Double mutations are extremely unlikely where the mutation rate is at or below the genome-wide average, so almost all will occur in mutation hotspots, and this is what I find. In all populations, the frequency of RHSVs per 100bp increases strikingly in the immediate vicinity of sites carrying three alleles. Crucially, the shape and width of the peak in RHSV number is very similar to equivalent plots based on sites that contribute most to D. The most parsimonious explanation is that both classes of site occur preferentially in narrowly define regions where the mutation rate is elevated. Other explanations appear unconvincing. For example, it could be argued that most triples arise through sequencing errors in regions where the alignments are problematic, thereby increasing the local frequency of sequencing errors. However, the similarity between plots for triples and plots of sites that contribute most to D are so similar that they surely share a common origin. Consequently, if the patterning around triples are dismissed as being due to sequencing errors signals used to infer introgression should also be dismissed as artefacts. In reality, I deliberately exclude singleton alleles from RHSV counts and so the contribution of sequencing errors to these patterns should be negligible.

It is interesting that the RHSV peak around sites where an ancestral archaic alleles is fixed in one population is noticeably stronger that for other classes of site, even after standardisation (see Figure 3). The reason for this is not immediately apparent. Under the inter-breeding hypothesis, it is difficult to see how the identity of the Denisovan base can exert any influence at all on the distribution of human-specific variants around introgressed fragments. However, under the variable mutation rate hypothesis, the recurrent mutations that drive D will differ depending on whether the Neanderthal allele is derived or ancestral. When derived, two independent A->B substitutions must occur, one in humans and one in Neanderthals. In contrast, when the Neanderthal allele is ancestral, a genuine back-mutation is often required (approximately half of all sites will become BBBA through a B->A substitution on the chimpanzee lineage, so would not involve a back-mutation in humans). Given the complexity of the processes that influence mutation type (SÉgurel *et al.* 2014; Harris 2015; Harris and Pritchard 2017) and rate (Conrad *et al.* 2011; Ségurel *et al.* 2014; Besenbacher *et al.* 2016; Sahakyan and Balasubramanian 2017), it is reasonable to expect back-mutations and parallel recurrent mutations to occur at different rates, but this aspect need more research.

With D being dominated by sites where ancestral archaic alleles are fixed in the population into which introgression is inferred and these sites lying in regions where mutation rate is unusually high in the other population, it is difficult to escape the conclusion that D is driven more by variation in rate of divergence from the ancestral state than by introgressed fragments. I have suggested before that a higher mutation rate in Africans could result from heterozygote instability (HI) (Amos 2010b; Amos 2013), where heterozygous sites become mismatches during synapsis that attract gene conversion events (Borts and Haber 1989; Borts *et al.* 1990; Collins and Newlon 1994; Nag *et al.* 1995). The resulting extra round of DNA replication provides additional opportunities for mutations to occur and will impact a region around 1-2Kb long, consistent with the way SNPs cluster (Amos 2010a). HI has the potential to drive mutation rate differences between populations wherever contrasting changes in heterozygosity have occurred. Humans provide and excellent test case, since around 25% of heterozygosity was lost during the out of African bottleneck (Prugnolle *et al.* 2005). Indeed, even Africans have a marginally lower mutation rat overall (Mallick *et al.* 2016) variants that likely post-date the out of Africa event reveal a substantially higher mutation rate in Africans compared with non-Africans, both as a genome-wide average and, across the genome, where relatively more heterozygosity has been lost (Amos 2013). It is also worth noting that striking mutation spectrum differences between human populations (Harris and Pritchard 2017) appear to be driven in part by a dependence on flanking heterozygosity (Amos 2019).

A model based on HI remains speculative but has a number of attractive features with respect to the patterns reported here. As discussed, back-mutations only become likely in regions where the local mutation is well above the genome-wide average. This is consistent with the observed association between D and difference in RHSV density. It is also consistent with the scale over which DHSV densities vary, which appear to be of the order of a few hundred bases, not dissimilar to size of gene conversion events (Borts and Haber 1989; Bertrán *et al.* 1997; Arbeithuber *et al.* 2015). Moreover, I have also found that flanking heterozygosity differences also correlate with D (Amos in review). Heterozygosity is only minimally impacted by RHSVs, particularly by the large number of RHSVs that are present in fewer than 10 copies across the entire dataset. Consequently, a correlation between D and local heterozygosity appears to require further explanation, and this could be provided by HI. HI should act to increase the mutation rate in regions with elevated heterozygosity, increasing both the frequency of RHSVs and the probability of BBBA sites acquiring back-mutations.

It could be argued that D is a rather crude measure of introgression and that more sophisticated methods should be preferred, including the inference of introgressed haplotypes (Patterson *et al.* 2012; Dannemann *et al.* 2016; Mallick *et al.* 2016; Peter 2016; Jacobs *et al.* 2019). However, while D is indeed rather a simple measure, both simulations and theory (Durand *et al.* 2011; Martin *et al.* 2015) show that it is effective and that it should capture any signal that is present. In this sense, evidence that the proportion of D attributable to introgressed fragments is small or even negligible cannot reasonably be negated by signals of introgression inferred by other methods. A more parsimonious explanation is that D and the inference of introgressed haplotypes are both failing to allow properly for variation in mutation rate and the accompanying higher than expected frequency of back-mutations. What is urgently needed is the development of methods that can infer introgressed fragments that are properly robust to back-mutations, variation in mutation rate to other aspects of the mutation process such as the way many mutations create clusters of up to eight substitutions on the same DNA strand (Besenbacher *et al.* 2016). Solving these issues may lead to a rather different view of hominin evolution.

## Methods

### Data

Data were downloaded from Phase 3 of the 1000 genomes project (1000 Genomes Project Consortium 2010) as composite vcf files from (ftp://ftp.1000genomes.ebi.ac.uk/vol1/ftp/release/20130502/). These comprise low coverage genome sequences for 2504 individuals drawn from 26 modern human populations spread across five geographic regions: Europe (GBR, FIN, CEU, IBS, TSI); East Asia (CHB, CHS, CDX, KHV, JPT); Central Southern Asia (GIH, STU, ITU, PJL, BEB); Africa (LWK, ESN, MSL, GWD, YRI, ASW, ACB); and the Americas (MXL, CLM, PUR, PEL). Individual chromosome vcf files for the Altai Neanderthal genome were downloaded from http://cdna.eva.mpg.de/neandertal/altai/AltaiNeandertal/VCF/. Vcf files for the Denisovan genome were downloaded from http://cdna.eva.mpg.de/denisova/VCF/human/. I focused only on homozygote archaic bases, accepting only those with 10 or more reads, fewer than 250 reads and where >80% were of one particular base. This approach sacrifices modest numbers of (usually uninformative) heterozygous sites but benefits from avoiding ambiguities caused by coercing low counts into genotypes.

### Data Analysis

Analysis of the 1000g data and archaic genomes were conducted using custom scripts to parse the data written in C++ (see Supplementary File 1 for an example script). The actual codes used for any given analysis are available on request. Sites were only considered if a base was called in all taxa: humans, the Altai Neanderthal, Denisovan and chimpanzee. Since the 1000 genomes data include much imputation, individual genotypes are likely less reliable than population frequencies. Consequently, D was always calculated probabilistically based on population allele frequencies and assuming independent assortment. Rare, human-specific variants (RHSVs) were defined as variants present as one of only two alleles in humans where the major allele is carried by the Neanderthal, Denisovan and chimpanzee, and where the minor allele frequency is greater 1 copy and less than 200 copies across the entire dataset of 2504 individuals. RHSVs were counted as the number of sites in a population that carry at least one copy of the rare variant.

Several different analyses involve counting RHSVs in 20 x 100bp windows around a central, focal base. Such analyses yield 20 RHSV counts for every pairwise combination of the 26 1000 genome populations (= 26 x 25 combinations). To make the resulting data more manageable, a number of averaging / standardisations were implemented. For RHSV counts, comparisons involving African populations yield much larger counts that other populations. For comparability, therefore, any given set of 20 sub-window counts were standardised by subtracting the 20-count average and then dividing by the same average, yielding values that reflect the proportional deviation from the mean. For the pairwise population comparisons, each involves an instance where one allele is fixed in one population and segregating in the other. Consequently, interest lies mostly with the allele that is present in only one population. To reflect this,

### Simulations

Simulated data were generated using the coalescent program ms (Hudson 2002). The model was coded:

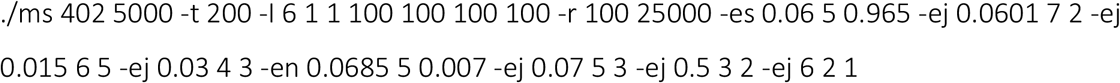

I assume a haploid population size of 10,000, a mutation rate of 10^−8^ and set theta to 200, such that each of 5,000 recombining fragments is 500Kb long. The hominin-chimpanzee split is taken to be 6,000,000 years ago, the Neanderthal-human split is here 500,000 years ago (generation length = 25 years. The human population splits into African and non-African lineages 70,000 years ago, the non-African lineage immediately experiencing a bottleneck that removes about 25% of heterozygosity. The African and non-African lineages each split into two at 30,000 and 15,000 years ago respectively. Introgression occurs at 60,000 years ago with a 3.5% injection into the non-African lineage.

## Acknowledgements

I thank Simon Martin, Rob Foley and Marta Lahr for many useful discussions.

